# Alternative Polyadenylation Regulates Patient-specific Tumor Growth by Individualizing the MicroRNA Target Site Landscape

**DOI:** 10.1101/601518

**Authors:** Soyeon Kim, Yulong Bai, Zhenjiang Fan, Brenda Diergaarde, George C. Tseng, Hyun Jung Park

## Abstract

**Background:** Alternative polyadenylation (APA) shortens or lengthens the 3’-untranslated region (3’-UTR) of hundreds of genes in cancer. While APA genes modify microRNA target sites in the 3’-UTRs to promote tumorigenesis, previous studies have focused on a subset of the modification landscape.

**Method:** For comprehensive understanding of the function of global APA events, we consider the total target site landscape of microRNAs that are significantly and collectively modified by global APA genes. To identify such microRNAs in spite of complex interactions between microRNAs and the APA genes, we developed <u>Pr</u>obabilistic <u>I</u>nference of <u>M</u>icroRN<u>A</u> <u>T</u>arget Site Modification through <u>APA</u> (PRIMATA-APA).

**Results:** Running PRIMATA-APA on TCGA breast cancer data, we identified that global APA events concentrate to modify target sites of particular microRNAs (<u>ta</u>rget-site-<u>mo</u>dified-<u>miRNA</u> or tamoMiRNA). TamoMiRNAs are enriched for microRNAs known to regulate cancer etiology and treatments. Also, their target genes are enriched in cancer-associated pathways, suggesting that APA modifies target sites of tamoMiRNAs to progress tumors. Knockdown of NUDT21, a master 3’-UTR regulator in HeLa cells, confirmed the causal role of tamoMiRNAs for tumor growth.

**Conclusions:** Further, the expressions of tamoMiRNA target genes, enriched in cancer-associated pathways, vary across tumor samples as a function of patient-specific APA events, suggesting that APA is a novel regulatory axis for interpatient tumor heterogeneity.

## Background

The dynamic usage of the messenger RNA 3’-untranslated region (3’-UTR) through alternative polyadenylation (APA) results in transcription of distinct isoforms with shortened or lengthened 3’-UTRs. 3’-UTR lengthening (3’UL) was recently reported to regulate cell senescence[1] linked with tumor suppressive pathways such as cell cycle inhibitors and DNA damage markers[2]–[5]. 3’-UTR shortening (3’US) was reported widespread in diverse types of human cancers[6]. Further, the impact of 3’US on prognosis[6] and its association with drug sensitivity[7] suggests clinical implications of APA events in human cancer.

We discovered a 3’US *trans* tumorigenic mechanism[8] based on its interaction with competing-endogenous RNA (ceRNA)[9]. Since 3’US removes microRNA (miRNA) target sites in the distal region of the 3’-UTRs, genes with 3’US event (3’US genes) do not sequester the miRNAs. Since ceRNAs co-regulate each other RNAs through competing for binding miRNAs, the miRNAs released from 3’US would be redirected to the ceRNA partners of the 3’US genes, working to promote tumorigenesis. Recently, we also found that this interactive mechanism between 3’US and ceRNA is involved in the subtype-specific progression of breast cancer[10]. However, computational predictions of ceRNA have been limited for RNAs with >5 miRNA target sites [8], [9], [11], [12]. Since such RNAs account for a subset of all expressed genes (e.g. among 17,085 genes with the average HiSeqV2 expression value > 1 across 1,098 TCGA breast tumor samples, 5,258 genes (30.5 %) have more than five microRNA binding sites), previous studies have not been able to bring comprehensive understanding of APA’s function. In this study, for each miRNA, we considered all expressed genes in which the target sites were modified by global APA events.

However, it is challenging to estimate the amount of APA-derived target site modification for miRNAs. First, while 3’US removes miRNA target sites in the 3’-UTRs, 3’UL plays a confounding role by adding the target sites back. Second, APA events and their associated miRNAs are on many-to-many relationships, making it difficult to pinpoint miRNAs that are collectively affected the most. To address these challenges, we developed a mathematical model that estimates the modification effect of global APA genes for each miRNA, <u>Pr</u>obabilistic <u>I</u>nference of <u>M</u>icroRN<u>A</u> <u>T</u>arget Site Modification through <u>APA</u> (PRIMATA-APA). We applied PRIMATA-APA on the TCGA breast cancer data[13] and found that APA modifies target sites of miRNAs known to regulate cancer etiology and treatments. Our prediction was further validated in the data from a previous study knocking down (KD) NUDT21, a master regulator of 3’-UTR shortening, in HeLa cell. Since the NUDT21 KD induced broad 3’UTR-shortening, leading to enhanced cellular proliferation in HeLa cells and glioblastoma tumor growth *in vitro* and *in vivo*, our validation confirms that APA events promote tumorigenesis by modifying target sites of miRNAs regulating tumorigenic process. Further analyses show that APA events affect tumor interpatient heterogeneity not dependent on miRNA expression changes but by modifying miRNA target sites.

## Methods

### TCGA breast tumor RNA-seq and miRNA-Seq data

Quantified gene expression files (RNASeqV1) for primary breast tumors and their matching solid normal samples were downloaded from TCGA Data Portal[51]. We used 106 breast tumor samples that have matched normal tissues. 10,868 expressed RefSeq genes (FPKM ≥ 1 in > 80% of all samples) were selected for downstream analyses. To better quantify gene expression in the presence of 3’-UTR shortening, we only used coding regions (CDS). Exon and CDS annotation for TCGA data and miRNA expressions (syn1445790) were downloaded from Sage Bionetworks’ Synapse database.

### Selection of miRNAs and genes

Predicted miRNA target sites were obtained from TargetScanHuman version 6.2[52]. Only those with a preferentially conserved targeting score (Pct) more than 0 were used[6]. Experimentally validated miRNA-target sites were obtained from TarBase version 5.0[53], miRecords version 4[54] and miRTarBase version 4.5[55]. The target sites found in indirect studies such as microarray experiments and high-throughput proteomics measurements were filtered out [56]. Another source is the microRNA target atlas composed of public AGO-CLIP data[57] with significant target sites (q-value < 0.05). The predicted and validated target site information was then combined to use in this study.

### <u>Pr</u>obabilistic <u>I</u>nference of <u>M</u>icroRN<u>A</u> <u>T</u>arget Site Modification through <u>APA</u> (PRIMATA-APA)

For transcript *x* with a constitutive proximal 3’-UTR (pUTR) and a distal 3’-UTR (dUTR), we previously defined the amount of target sites for miRNA miR_j_ in all copies of transcript *x* as follows[8].

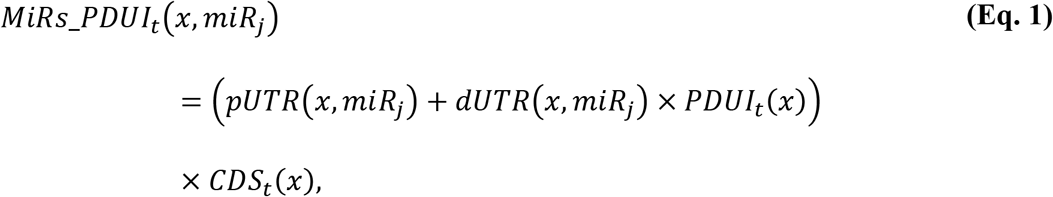

where *pUTR(x, miR_j_)* and *dUTR(x, miR_j_)* are the numbers of miR_j_ target sites in pUTR and dUTR of x. *PDUI_t_(x)* is the Percentage of dUTR Usage Index[6] of x and *CDS_t_(x)* is the expression level of x in a tumor sample. Note that *MiRs_PDUI_n_(x, miR_j_)* can be calculated for a normal sample with *PDUI_n_(x)* and *CDS_n_(x)*. If APA-derived miRNA target site modification is not considered, the amount of target sites for *miR_j_* in all copies of transcript *x* would be calculated as follows:

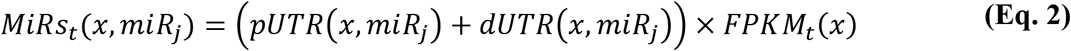

Based on **Eq.1** and **Eq.2**, PRIMATA-APA calculates *MiRs_PDUI_t_(miR_j_)* and *MiRs_t_(miR_j_)* defined as below.

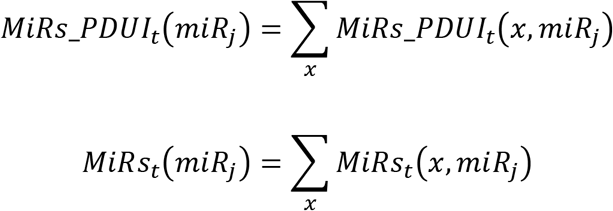

With *MiRs_PDUI_t_*(*miR_j_*), *MiRs_t_*(*miR_j_*), *MiRs_PDUI_n_*(*miR_j_*), and *MiRs_n_*(*miR_j_*) in a contingency table, PRIMATA-APA estimates significance of target site modifications for *miR_j_* by testing nonrandom association in tumor and normal states (using χ^2^ test), followed by FDR control using FowardStop[58] (FDR < 0.01).

## Results

### Collective impact of APA genes for the *trans* effect

To identify APA events in large-scale data, several computational tools have been developed that use RNA-Seq data[6], [14]–[17]. For example, statistically significant APA genes can be defined using the difference in Percentage of Distal polyA site Usage Index (ΔPDUI)[6]. Among the significant APA genes from such tools, current analyses have focused mostly on a subset of them that strongly changed 3’-UTR usage in tumor (strong APA genes). For example, studies on TCGA human cancer[6] and cell line data[7] focused on the strong APA genes (ΔPDUI < −0.2 for 3’US and ΔPDUI > 0.2 for 3’UL) selected from significant APA genes (FDR < 0.05). However, strong APA genes account for only a small portion of all significant APA genes. For example, in TCGA tumor-normal sample pair BH-A1FJ with the greatest number of significant APA genes, the strong APA genes account only for 50.5% (1,523) of 3,015 significant APA genes (**Fig. 1A**). Across 106 breast tumor-matched normal sample pairs in TCGA, this trend is more pronounced in that only 40.5% of significant APA genes are strong APA genes on average (**Fig. 1B**).

**Figure 1.**
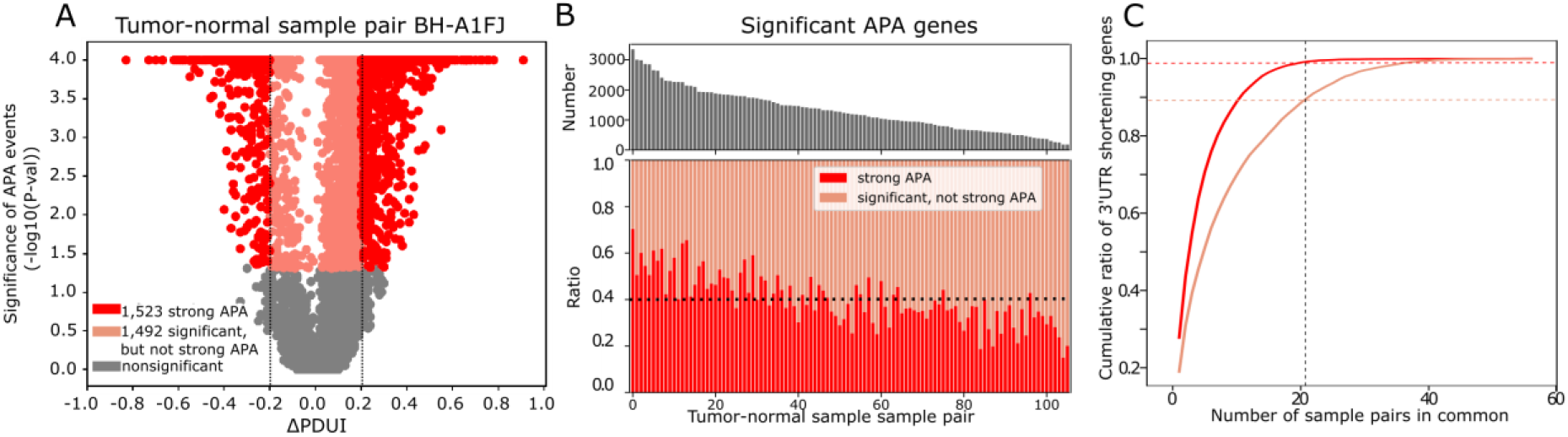
Collective impact of strong and significant APA events. A. Statistical significance of APA genes in a breast tumor-normal sample pair (TCGA-BH-A1FJ) with their ΔPDUI (Percentage of Distal polyA site Usage Index) values (tumor-normal). Since PDUI represents the ratio of isoforms with distal 3’-UTR, negative ΔPDUI value represent 3’-UTR shortening target genes and positive ΔPDUI value 3’UL genes. Strong APA target genes are in red, significant-but-not-strong ones in pink and not significant ones in gray. B. For 106 breast tumor-normal sample pairs sorted by the number of significant APA target sites, upper panel shows the total number of significant APA genes and the lower panel shows the ratio of the APA genes by whether it is significant-but-not-strong (orange) or strong (red). Black dotted line represent the average ratio of strong APA genes. C. Cumulative ratio of 3’US genes shared by sample pairs. Red and orange dotted lines represent the ratio of strong and significant 3’US genes shared by < 21 sample pairs, respectively.

We found that significant APA genes, strong or not, together elucidate the common *trans* mechanism of APA. In 106 normal/tumor sample pairs, we identified totally 6,825 significant 3’US genes, of which 82.4% (5,626) are strong (ΔPDUI < −0.2) in some other sample pairs, demonstrating that significant 3’US genes in a sample pair are likely to undergo a strong 3’-UTR shortening in other sample pairs. Further, considering all significant 3’US genes identifies much more recurring 3’US genes while increasing only 15.9% (6,825 vs. 5,626) more 3’US genes in total. For example, 5.5-fold more 3’US genes recur across 20% (21) of the 106 normal/tumor sample pairs if all significant 3’US genes are considered (613 significant vs. 110 strong only, **Fig. 1C**). In the previous studies studying cis effect of APA events, focusing on strong APA genes would be reasonable, since strong 3’US genes are likely strong *cis* targets of 3’US. However, as 3’US events exert the *trans* mechanism through modifying miRNA target sites[8], significant-but-not-strong 3’US genes contribute to the *trans* mechanism by modifying miRNA target sites on their 3’UTRs[18], especially if they are highly expressed.

In the same sense, 3’UL genes need to be considered, since they would increase miRNA target sites to compensate for those decreased by 3’US genes. Previous studies reported smaller numbers of 3’UL genes than 3’US from TCGA cancer patients[6] or cancer cell lines[7]. This was partly because they focused only on strong APA genes common to the samples. Considering all significant APA genes in our TCGA breast cancer analysis, we found that 3’UL is as widespread as 3’US. For example, in TCGA tumor-normal sample pair AC-A2FB, which has the smallest ratio of strong APA genes to significant APA genes, 90.1% of the significant APA genes are 3’UL genes (**S. Fig 1A**). In each of 106 TCGA breast tumor-matched normal sample pairs, average 56.4% of significant APA genes are 3’UL genes (**S. Fig. 1B**). As in the case of 3’US, considering significant 3’UL genes enables to identify much more recurring 3’UL genes than considering only strong 3’UL genes. In our TCGA breast cancer data, 83.7% of the total significant 3’UL genes (6,081 of 7,265) are strong (ΔPDUI > 0.2) in some other samples and 3.5-fold more 3’UL genes recur in 20% of the sample pairs when all significant 3’UL genes are considered (993 significant vs. 288 strong only, **S. Fig. 1C**). Based on the results, investigating the common *trans* mechanisms of APA requires to consider all significant APA genes of both 3’UL and 3’US.

### Probabilistic Inference of MicroRNA Target Site Modification through APA (PRIMATA-APA)

To investigate the common *trans* mechanisms of APA that modifies miRNA target sites, we quantify target site modifications due to significant APA events for each miRNA by developing a mathematical model, <u>Pr</u>obabilistic <u>I</u>nference of <u>M</u>icroRN<u>A</u> <u>T</u>arget Site Modification through <u>APA</u> (PRIMATA-APA). Previously, we successfully predicted gene expression changes based on the estimated number of miRNA target sites in the presence of 3’US (*MiRs_PDUI_t_*(*x, miR_j_*), **Eq. 2**)[8]. By extending this estimation, PRIMATA-APA estimates the total number of target sites for each miRNA with and without consideration of APA events, 3’US and 3’UL together. Based on the difference of the estimations, PRIMATA-APA evaluates the statistical significance of the APA-derived modification for each miRNA with the direction of the modification, either increase or decrease (see Methods). Since miRNAs play roles mainly by binding to the target sites of genes [19], considering miRNA target site information should help us to understand their effect on transcriptomic dynamics beyond miRNA expression information.

### Global miRNA target site modification due to alternative polyadenylation

To identify miRNAs affected by APA-derived target site modification, we ran PRIMATA-APA on the 70 breast tumor-matched normal samples out of 106 pairs for which miRNA expression information was available. In the data, we considered 588 moderately expressed miRNAs (> 1 and < 100 FPM on average) on the expressed genes (avg. FPKM > 1) in each sample pairs. In 39 (55.7%) of the 70 breast tumor samples, PRIMATA-APA identifies significant target site modifications (FDR < 0.01) for more than 100 miRNAs (**Fig. 2A, B**). Further, in each sample pair, miRNA target sites are either mostly increased or decreased, which makes a negative correlation between the number of target sites increased and decreased across tumor-normal sample pairs (P=0.006, **Fig. 2C**).

**Figure 2.**
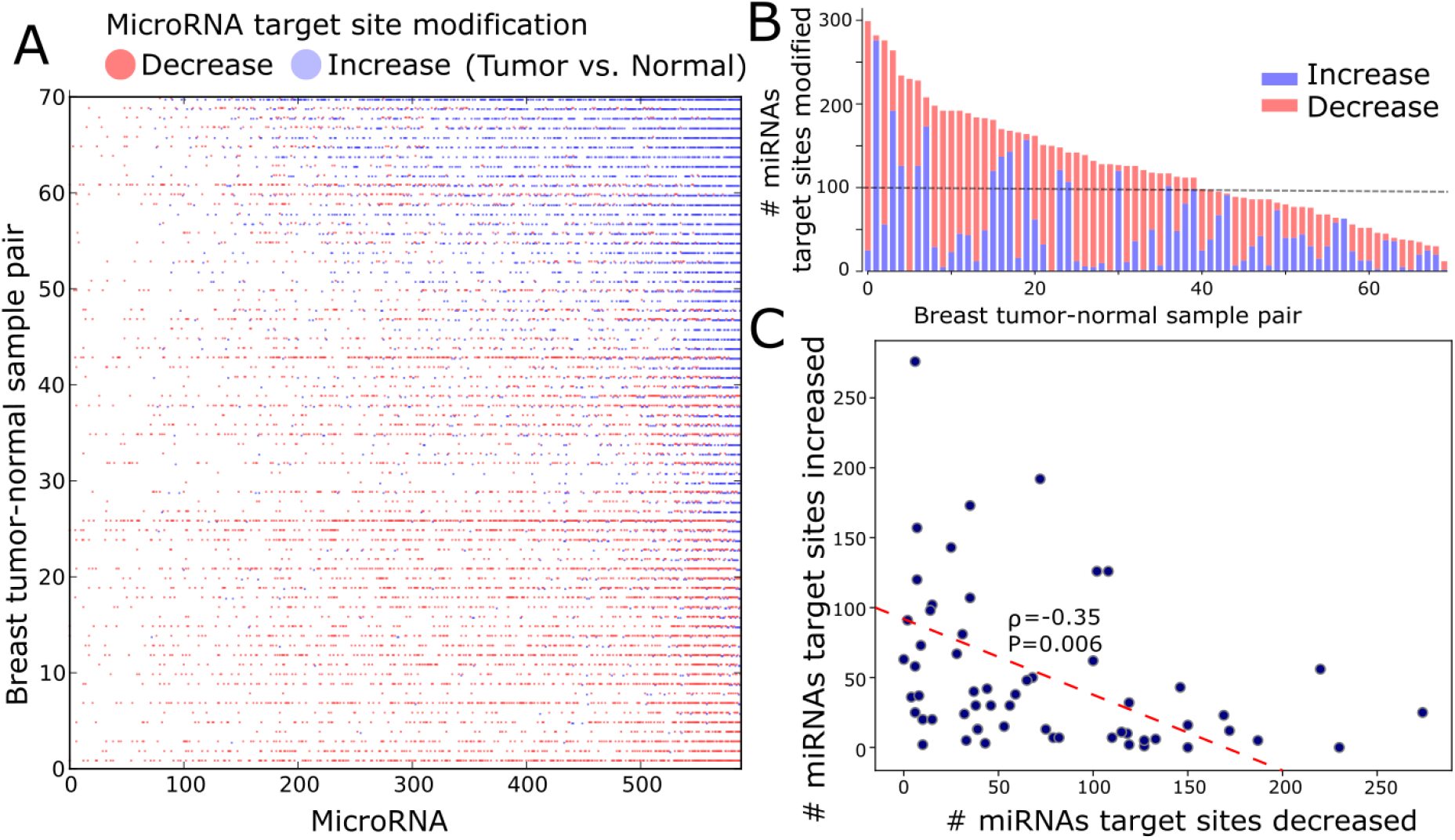
Tumor-specific APA-derived microRNA binding site changes. A. The heatmap shows tumor-normal samples (row) where the total number of binding sites for each microRNA (column) is increased (blue) or decreased (red) due to APA. Not significant changes or no changes are not colored. Samples are sorted by the number of increased microRNA target site modicifcation. B. The total number of miRNA binding site changes, either increased (blue) or decreased (red) due to APA, in breast tumor-normal samples pair sorted by the total number of modification per sample pair. C. Number of miRNAs of which binding sites are increased (y-axis) or decreased (x-axis) in each tumor-normal sample. The red dotted line represents linear least-squares regression.

Additionally, we found that APA events modify miRNA target sites in a subtype-specific manner. The five subtypes of breast cancer by PAM50 are known to involve distinct molecular pathways with different clinical outcomes[20]. In our TCGA breast cancer data, none of basal and Her2 subtype samples increases target sites for > 100 miRNAs, while 57.1% (4/7) of the samples decrease target sites for > 100 miRNAs, indicating a significant (P=0.009) bias toward miRNA target site decrease. However, other PAM50 subtypes (Luminal A, B, and Normal-like) do not show such a bias (**S. Fig. 2**). Since both basal and Her2 subtypes are close in terms of molecular pathways and worse prognosis (reviewed in [21], [22]), their common pattern in miRNA target site modification suggests a similar APA landscapes between them. Altogether, the results show that APA globally modifies miRNA target sites for breast cancer in a non-random and a subtype-specific manner.

### APA modifies target sites of miRNAs associated with cancer

To perform subsequent analyses with the focus on particular miRNAs, we identified miRNAs whose target sites are more likely modified by APA events. Based on the number of tumor samples where the target sites are significantly increased or decreased, we selected top half (289) of 588 moderately expressed miRNAs, whose target sites are more often modified than the other (299) miRNAs, which will be termed <u>ta</u>rget site <u>mo</u>dified miRNA (tamoMiRNA). APA events modify target sites of tamoMiRNAs significantly more than the other miRNAs (P-value=1.12e^-28^, **Fig. 3A**). The 289 tamoMiRNAs are significantly (P=5.8e^-5^) enriched in the miRNA sets known for cancer etiology and treatments compared to the other miRNAs. Specifically, tamoMiRNAs are enriched in the miRNAs that are dysregulated in breast cancer with clinical and biological implications[23], regulating diverse mechanisms for breast cancer[24], regulatory elements in either adaptive or innate immune system[25], or potential prognostic and predictive biomarkers identified for breast cancer[26](**Fig. 3B**, S. Table 1). Among 43 tamoMiRNAs found in the categories, 31 (72.1%) occur only in one of the categories (**S. Fig. 3**), confirming that the high enrichment of tamoMiRNAs to the multiple categories reflects their important roles in tumor, not redundancy in data curation. Also, we estimated conservation score (PhyloP[27], 46 way Placental) of 202 tamoMiRNAs and 191 other miRNAs for which miRBase[28] uniquely curated the genomic locations. TamoMiRNAs have significantly (P= 7.19e^-5^) larger conservation scores than the other miRNAs (**Fig. 3C**), implicating their functional significance. Altogether, these results indicate that, by selection or design, APA modifies target sites of miRNAs that are evolutionary conserved and functionally important for cancer etiology and treatments.

**Figure 3.**
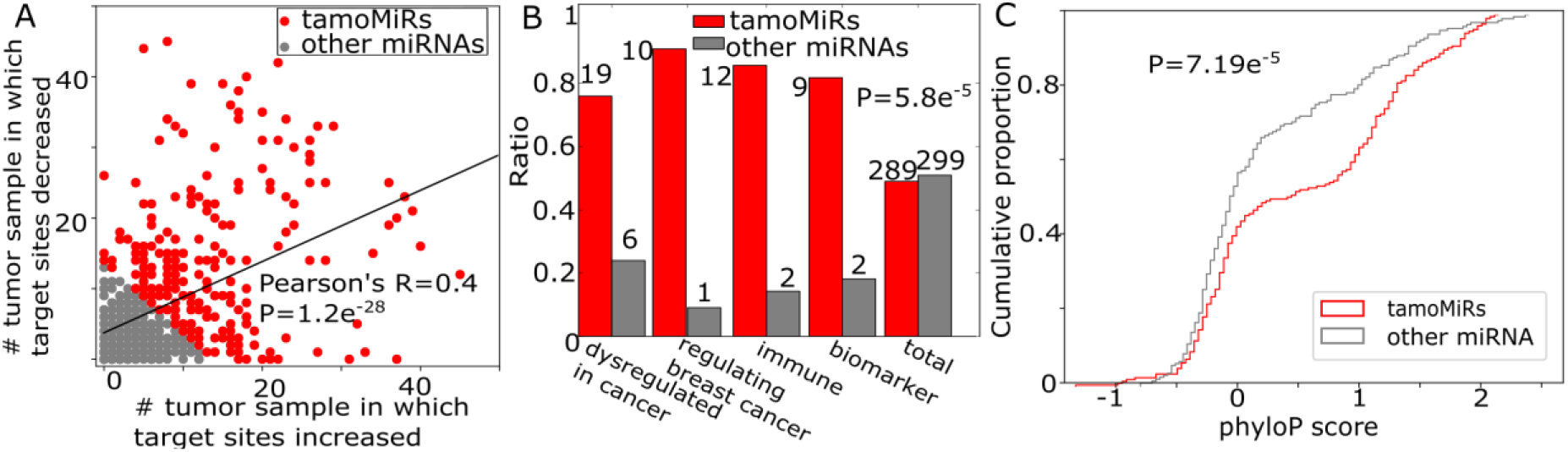
APA modifies binding sites of miRNAs associated with cancer. A. The number of tumor-normal samples between which binding sites for each miRNA are increased (x-axis) or decreased (y-axis). For further analyses, we dichotomize miRNAs by the amount of binding site modification into tamo- (red) and the other (gray) miRNAs. B. Number of cancer-related miRNAs in tamo- (red) and the other (gray) miRNAs. C. Distribution of phyloP conservation score for 202 tamo- and 191 the other miRNAs.

### APA promotes tumorigenesis by modifying target sites of miRNAs regulating cancer

To gain insights into the cause-and-effect relationship between tamoMiRNAs and tumor growth, we revisited a previous study[29], in which NUDT21 knockdown (KD) induced broad 3’UTR-shortening, leading to enhanced cellular proliferation in HeLa cells and glioblastoma tumor growth *in vitro* and *in vivo*. Our analyses on NUDT21 experiment data (available in GEO under GSE42420 and GSE78198) validated our observations above. First, both significant-but-not-strong and 3’UL genes need to be considered for a comprehensive understanding of APA’s function. Since NUDT21 is a master regulator for 3’UTR-shortening, knocking down NUDT21 yields much more strong 3’US genes (2,550 genes with ΔPDUI < −0.2 and FDR < 0.05) than 3’UL genes (84 with ΔPDUI > 0.2 and FDR < 0.05) in HeLa cells (**Fig. 4A**). However, this experiment also yielded many significant-but-not-strong APA genes (615 3’UL vs. 2,847 3’US) that modifies miRNA target site landscape, where a substantial ratio (17.7%) are 3’UL genes. Second, miRNA target site modification due to APA genes is a robust estimator for the *trans* effect of APA genes. To show this, we performed the following analysis on each of 383 miRNAs whose expression was available in the experiment data[8]. Given two replicate data of wild type HeLa cells (WT) and NUDT21 KD, we paired a WT and a NUDT21 KD data to make two pairs. Then, we ran PRIMATA-APA for each miRNA to see whether the target sites increase or decrease in each pair (NUDT21 KD vs. WT). Between the pairs, we validated that the PRIMATA-APA analysis results agree across the miRNAs using Cohen’s κ (P < 10e^-16^) with 97.4% of agreement (using R package ‘irr’). For further analyses, we identified tamoMiRNAs for NUDT21 KD experiment as follows. With the high agreement of PRIMATA-APA calls between the sample pairs, we ran PRIMATA-APA on the averaged expression data for two WTs and two NUDT21 KD cells (S. Table 2). Then, we ranked the miRNAs based on the χ^2^ value that indicates the degree of APA-derived target site modification (see Methods). Then, as before, the top half of the miRNAs (191) are tamoMiRNAs for NUDT21 KD experiment. It is interesting that miR-3187-3p that was extensively validated to repress tumor suppressor genes for tumorigenesis in the NUDT21 KD experiment is a tamoMiRNA[8]. We found that this experiment shares not only a significant number of strong and significant 3’US genes (91, P=9.3×10^-21^, **Fig. 4B**), but also a significant number of tamoMiRNAs (119, P=1.16×10^-11^, **Fig. 4C**) with the TCGA breast cancer analysis. Since both the TCGA data (tumor vs. normal) and NUDT21 KD data represent a tumorigenic process based on 3’US, these overlaps indicate that 3’US genes and miRNA target sites they modify contribute to the tumorigenic process. Furthermore, as the case of TCGA breast cancer data, tamoMiRNAs for NUDT21 KD are highly enriched for miRNA sets known for cancer etiology and treatments (P=0.00014, **Fig. 4D**) and evolutionary conserved (P=0.04, **Fig. 4E)**, confirming the functional importance of tamoMiRNAs. NUDT21 KD validated the cause-and-effect relationship from APA events to tumor growth through modifying target sites of tamoMiRNAs.

**Figure 4.**
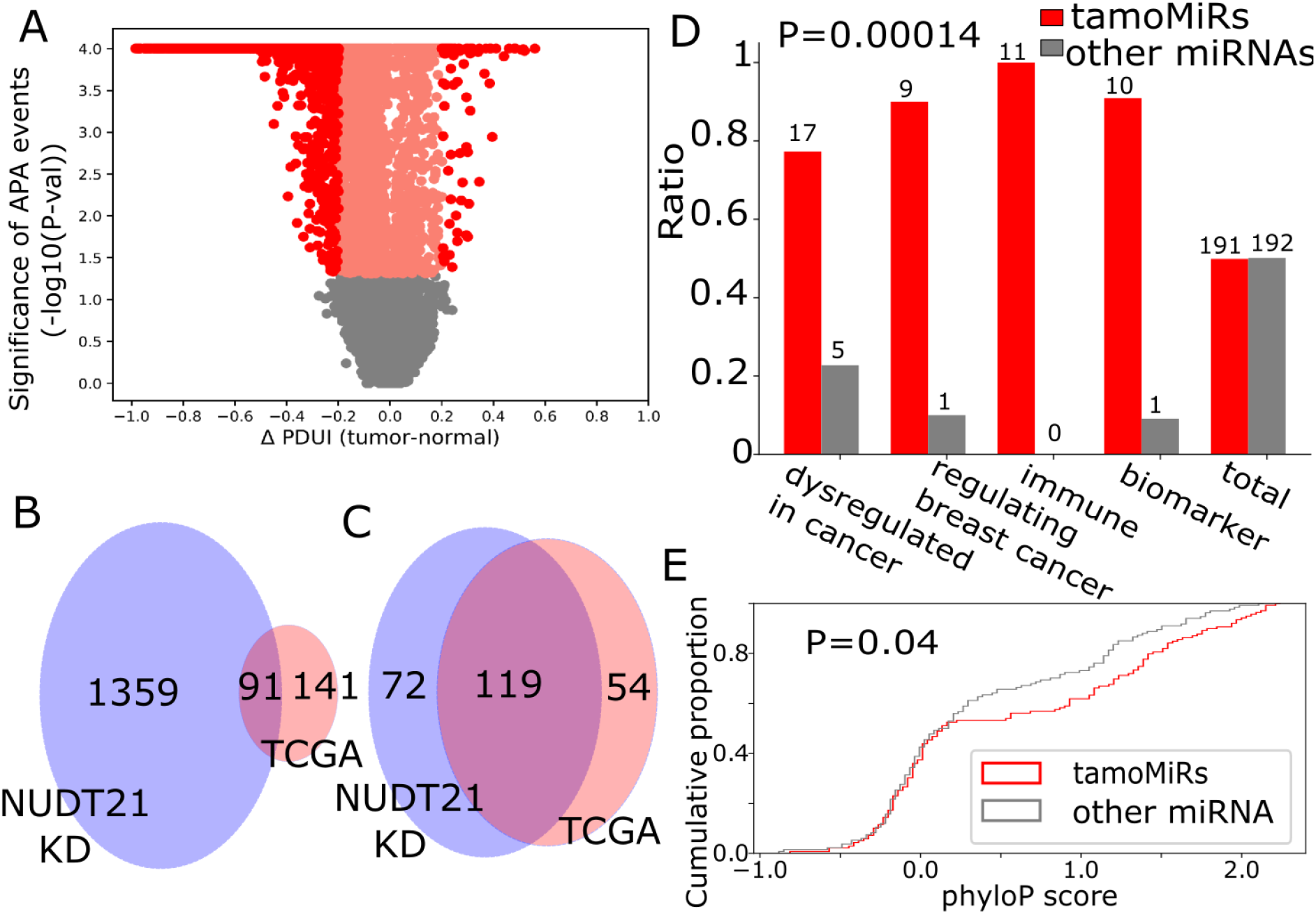
APA promotes tumorigenesis by modifying target sites of miRNAs regulating cancer. A. Statistical significance of APA genes in NUDT21 KD experiment data with their ΔPDUI values (tumor-normal). Overlap of B. 3’US genes and C. tamoMiRNAs between TCGA breast cancer data and NUDT21 KD data based on PRIMATA-APA. D. Number of cancer-related miRNAs in 191 tamo- (red) and the other (gray) 192 miRNAs. E. Distribution of phyloP conservation score for 139 tamo- and the other 134 miRNAs.

### APA modifies target sites of miRNAs to effectively regulate biological processes

To further investigate the function of the miRNA target site modification in TCGA breast cancer data, we focused on the target genes of the tamoMiRNAs (see Methods). First, we identified GO terms enriched for tamoMiRs using MiEAA[30]. Probably due to the many-to-many relationships between miRNAs and target genes[31]–[33], inputting all tamoMiRNAs and all the other miRNAs to MiEAA web server returns mostly under-represented terms. So, we focus MiEAA analysis on 99 tamoMiRNAs (with the greatest number of samples in which modified) and 105 other miRNAs (with the least number of samples in which modified). 125 and 1 biological terms are significantly (FDR < 0.01) enriched for tamoMiRNAs and for the other miRNAs respectively (S. Table 3). The significant bias (P-value=5.0×10^-5^) of the number of enriched biological terms to tamoMiRNAs suggests that APA events effectively regulate biological functions. Additionally, compared to the other miRNAs, tamoMiRNAs are exclusively enriched for pathways with keyword “growth factor”, “signaling”, and “circadian”, (**Fig. 5A**, S. Table 3), which are essential for tumor initiation and progression[34].

**Figure 5.**
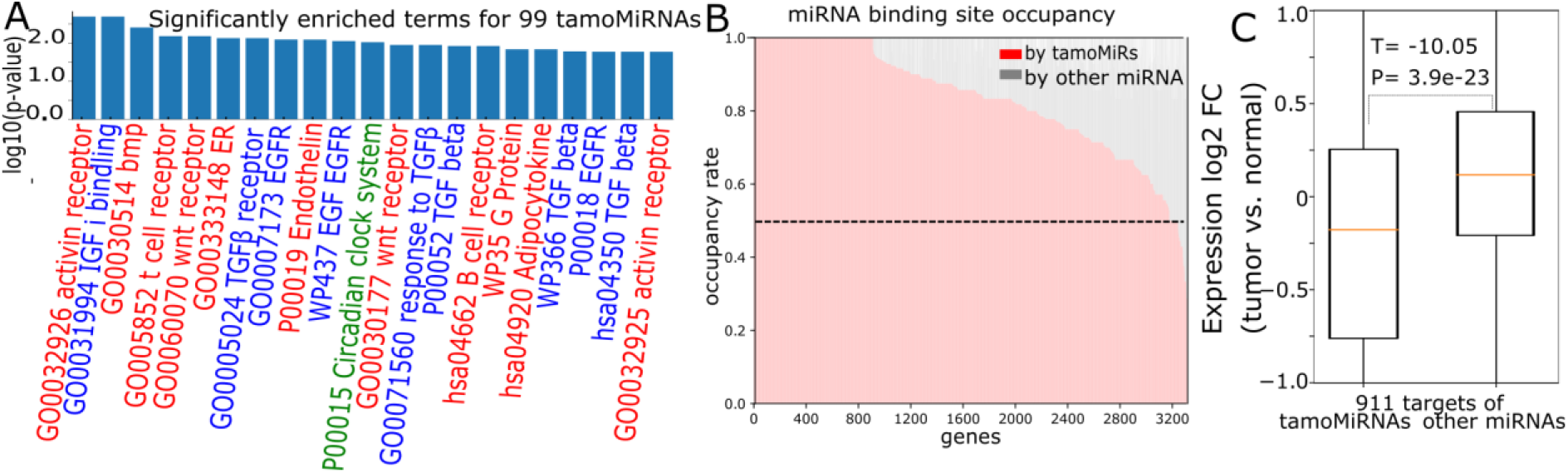
TamoMiRNAs effectively regulate biological processes. A. Cancer-associated pathways enriched for 99 tamoMiRNAs with their enrichment p-values (red for “signaling”, blue for “GF” (growth factor), and green for “circadian”). B. Number of target sites for tamoMiRs and the other miRNAs in the genes with more than 5 target sites. C. Expression fold change (log2 tumor vs. normal) of 911 genes that are targets of tamoMiRs and other miRNAs.

To understand their impact in regulating gene expression, we evaluated the number of genes targeted by tamoMiRNAs. Among 3,318 expressed genes in the breast tumor data that are likely controlled by miRNAs (> 5 miRNA target sites), 3,177 genes (95.7%) have more target sites for tamoMiRNAs than for the other miRNAs (**Fig. 5B**). Further, 911 of 3,177 (27.4%) genes have target sites only for tamoMiRNAs in their 3’-UTRs. While expression fold change (tumor vs. normal) does not differ between tamoMiRNAs and other miRNAs (P=0.1, **S. Fig. 2**), 911 genes targeted only by tamoMiRNAs are significantly more down-regulated in tumor (P=3.9e^-23^) than the same number of genes affected by other miRNAs (**Fig. 5C**), suggesting that tamoMiRNAs effectively regulate gene expressions of target genes in tumor. Altogether, the results indicate that APA modifies target sites of miRNAs that effectively regulate genes in tumor-associated pathways.

### APA regulates tumor-specific progression to affect interpatient heterogeneity

As miRNAs are known to regulate cancer interpatient heterogeneity[35], we further studied the effect of APA-derived miRNA target site modification on cancer interpatient heterogeneity. In particular, since 3’-UTR shortening regulates gene expression by modifying miRNA target sites[8], we hypothesized that variation in the degree of APA events across tumor samples diversifies the effect of miRNAs on the target genes for tumor interpatient heterogeneity. To test this hypothesis, we compared the expression variation of tamoMiRNAs, the other miRNAs and 911 of their target genes defined above. While the expression variation across the sample pairs changes equally in tumor for both tamoMiRNAs and the other miRNAs (P=0.4, **Fig. 6A**), the expression variation of genes that are targeted by tamoMiRNA is significantly higher than that of the other miRNAs (P=4.9e^-15^, **Fig. 6B**) in tumors. Since the degree of APA events varies significantly more (P=0.004) in tumor (**S. Fig. 6B**), the high variation in tamoMiRNA target gene expression in tumor is, at least partly, attributable to the APA events modifying tamoMiRNA target sites in the genes.

**Figure 6.**
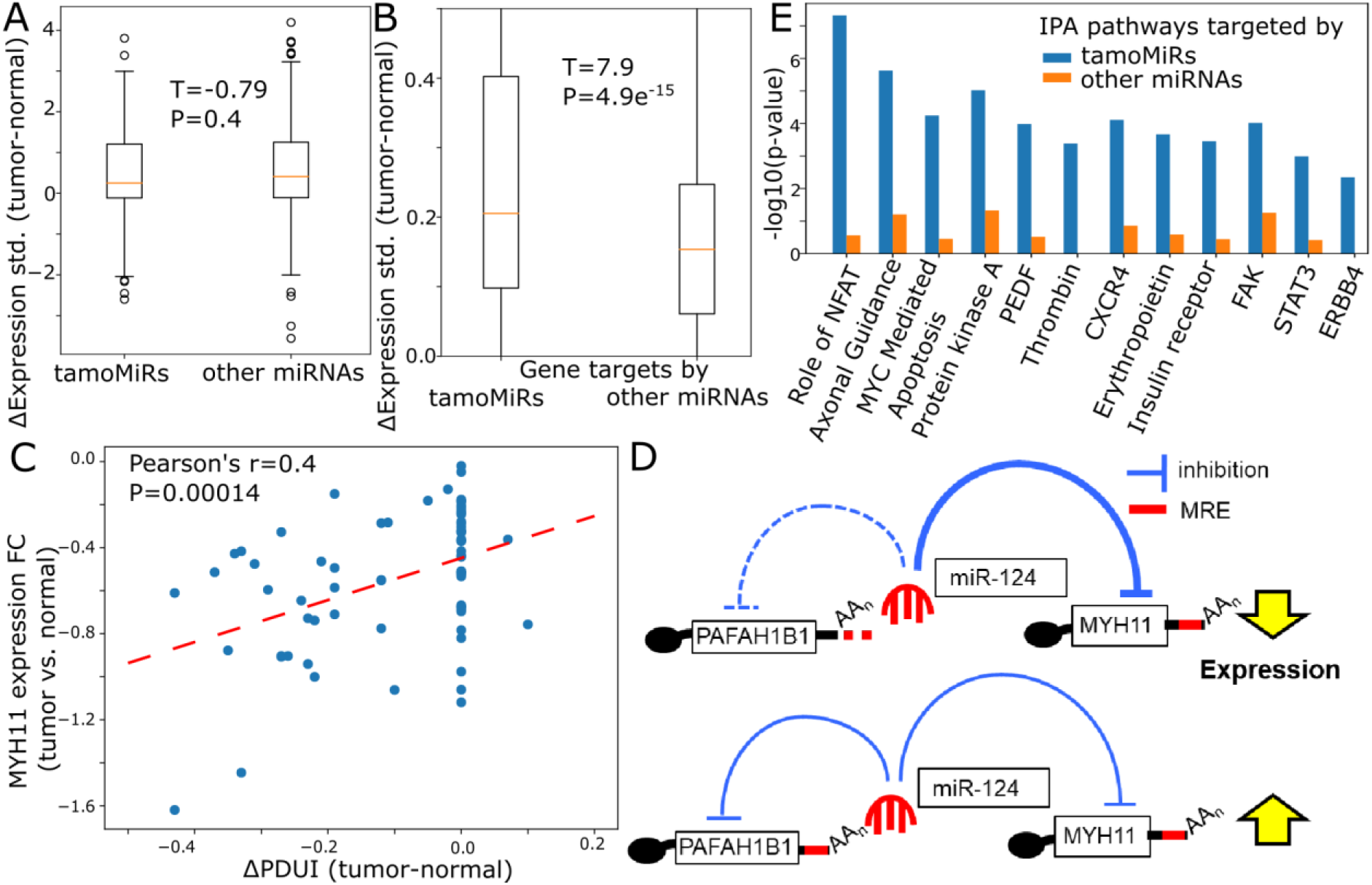
APA regulates tumor-specific progression in trans. Distribution of standard deviation values across sample pairs on the expression values of A. tamoMiRs and other miRNAs and B. their target genes. C. Scatterplot of MYH11 expression fold change values against its ΔPDUI values. The red dotted line represents linear least-squares regression. D. Illustration of the effect of PAFAH1B1’s 3’-UTR shortening on MYH11 expression mediated by miR-124/124ab/506. MRE stands for microRNA recognition element. E. IPA comparison analysis between gene targets by highly and lowly modified miRNAs for pathways implied for cancer progression and migration, NFAT[59], Axonal Guidance[60], MYC Mediated Apoptosis[61], Protein kinase A[44], Pigment epithelium-derived factor (PEDF)[62], Thrombin[63], CXCR4[43], Erythropoietin[64], Insulin receptor[65], FAK[66], STAT3[67].

An example is myocin heavy chain 11 (MYH11) disruption of which is associated tumorigenesis of various types of cancer [36]–[40]. According to our miRNA-target site information, MYH11 is predicted to have a target site for a single miRNA (miRNA-124/124ab/506). Although the expression change of MYH11 is not correlated with that of the miRNA expression (**S. Fig. 5A**), it is significantly (Pearson’s r=0.4, P-value=0.00014, **Fig. 6C**) correlated with ΔPDUI values of its ceRNA partner gene, platelet-activating factor acetylhydrolase IB subunit alpha (PAFAH1B1). The functional role of PAFAH1B1 in tumor has not been widely studied, especially in association with MYH11. Our analysis found that 3’-UTR of PAFAH1B1 undergoes a significant (T-test statistic=2.65, P=0.004) shortening in tumor vs. normal. Since the 3’-UTR shortening of PAFAH1B1 in tumor (negative ΔPDUI values [6]) is expected to repress MYH11 (**Fig. 6D**), the positive correlation between MYH11 expression and ΔPDUI of PAFAH1B1 (**Fig. 6C**) supports the role of 3’US *trans* effect differentiating MYH11 expression. It is worth noting that the mediating miRNA, miRNA-124/124ab/506, is a tamoMiRNA (S. Table 1).

Further enrichment analyses support that APA-derived, miRNA-mediated[41], [42] transcriptomic diversity contributes to interpatient difference in tumor progression. First, 911 genes targeted only by tamoMiRNAs include significantly more oncogenes than the same number of genes targeted by other miRNAs (43 vs. 28, P-value=0.03), suggesting that a varying degree of APA diversifies the effect of miRNA target activity on the oncogenic processes across tumor samples. Second, Ingenuity Pathway Analysis showed that cancer progression and migration pathways are more enriched in tamoMiRNA target genes than the target genes of the other miRNAs (P-value < 10^-3^, **Fig. 6E**, S. Table 4). Specifically, they are implicated for breast cancer often through miRNAs, e.g. with miR-494 suppressing chemokine (C-X-C motif) receptor 4 (CXCR4) for breast cancer progression[43], miR-200c regulating Protein kinase A subunits for cancer cell migration[44], and miR-520b targeting Interleukin-8 for breast cancer cell migration[45] (see other examples in **Fig. 6E**). Since these miRNAs (miR-494, miR-200c, and miR-520b) are tamoMiRNAs of which APA modifies target sites, our results suggest that APA diversifies miRNA target site landscape for tumor-specific progression.

## Discussion

Here, we studied the post-transcriptional regulation efficiency of miRNAs in presence of APA events. As miRNAs bind and repress their target genes in the post-transcriptional level, a comprehensive understanding of miRNAs’ function should be based not only on their expression levels but also on the target site landscape. However, previous computational works predicted miRNAs’ function based mostly on miRNA expression changes. For example, miRNAs up-regulated in tumor were considered to be oncogenic miRNAs and those down-regulated were tumor-suppressive [46], [47]. On the other hand, we characterized miRNA target site landscape modified by APA events using a mathematical model, PRIMATA-APA. Further analyses on TCGA breast cancer data clearly demonstrated that considering miRNA target site modification brings a comprehensive understanding of miRNA function in the presence of widespread APA events in tumor.

Further, our work sheds novel insights into the development of therapeutic miRNAs. The high enrichment of tamoMiRNAs to miRNAs known to regulate cancer (**Fig. 3B**) suggest that APA events that are associated with tumorigenesis[8], prognosis[6] and treatment outcomes[48] make the associations by modifying target sites of tamoMiRNAs. Based on this rationale, further inspection of tamoMiRNAs would effectively narrow down miRNA search space to identify novel therapeutic miRNAs. Further, identifying miRNAs significantly associated with interpatient heterogeneity will help regulate interpatient tumor heterogeneity, which is essential for the success of early cancer detection and the development of new effective therapies[49], [50]. For example, since MYH11’s oncogenic effect is associated with a varying degree of APA events of PAFAH1B1 through miRNA-124/124ab/506 in breast cancer, molecular agents for miRNA-124/124ab/506 may help normalize different MYH11 effect on cancer patients.

## Declarations

**Ethics approval and consent to participate**

### Consent for publication

This study does not involve human participants.

### Availability of data and materials

The open source PRIMATA-APA program (version 0.9.2) is freely available at https://github.com/thejustpark/PRIMATA-APA/ with necessary example data for this analysis.

### Author Contributions

H.J.P and S.K. conceived the project, designed the experiments and performed the data analysis. Y.B. and Z.F. performed the subtype analysis. H.J.P and S.K. wrote the manuscript with input from B.D., G.C.T..

## Acknowledgements

This research was supported in part by the University of Pittsburgh Center for Research Computing through the resources provided.

---

**S. Figure 1.**
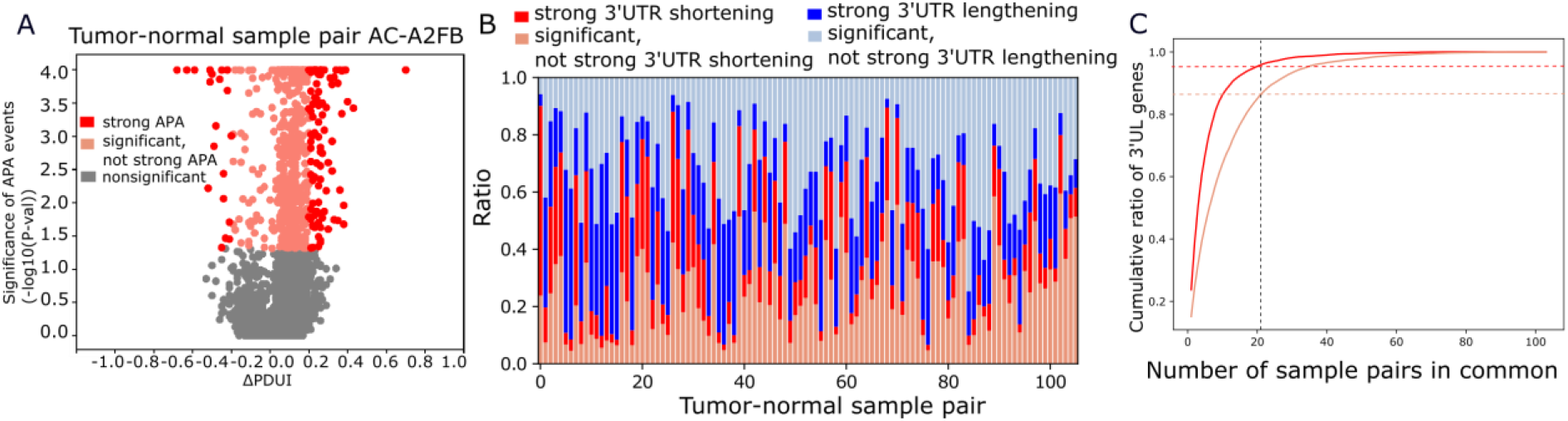
A. Statistical significance of APA target genes in a breast tumor-normal sample pair (TCGA-BH-A1FJ) with their ΔPDUI (Percentage of Distal polyA site Usage Index) values (tumor-normal). B. In each of 106 breast tumor-normal sample pairs, the ratio of the APA target genes by whether it is significant and not strong or strong, also by whether it is 3’UTR shortening or lengthening. They are ordered in consistency with Fig. 1B. C. Cumulative ratio of genes with lengthened 3’-UTRs shared by sample pairs. Cumulative ratio of 3’US genes shared by sample pairs. Red and orange dotted lines represent the ratio of strong and significant 3’US genes shared by < 21 sample pairs, respectively.

**S. Figure 2.**
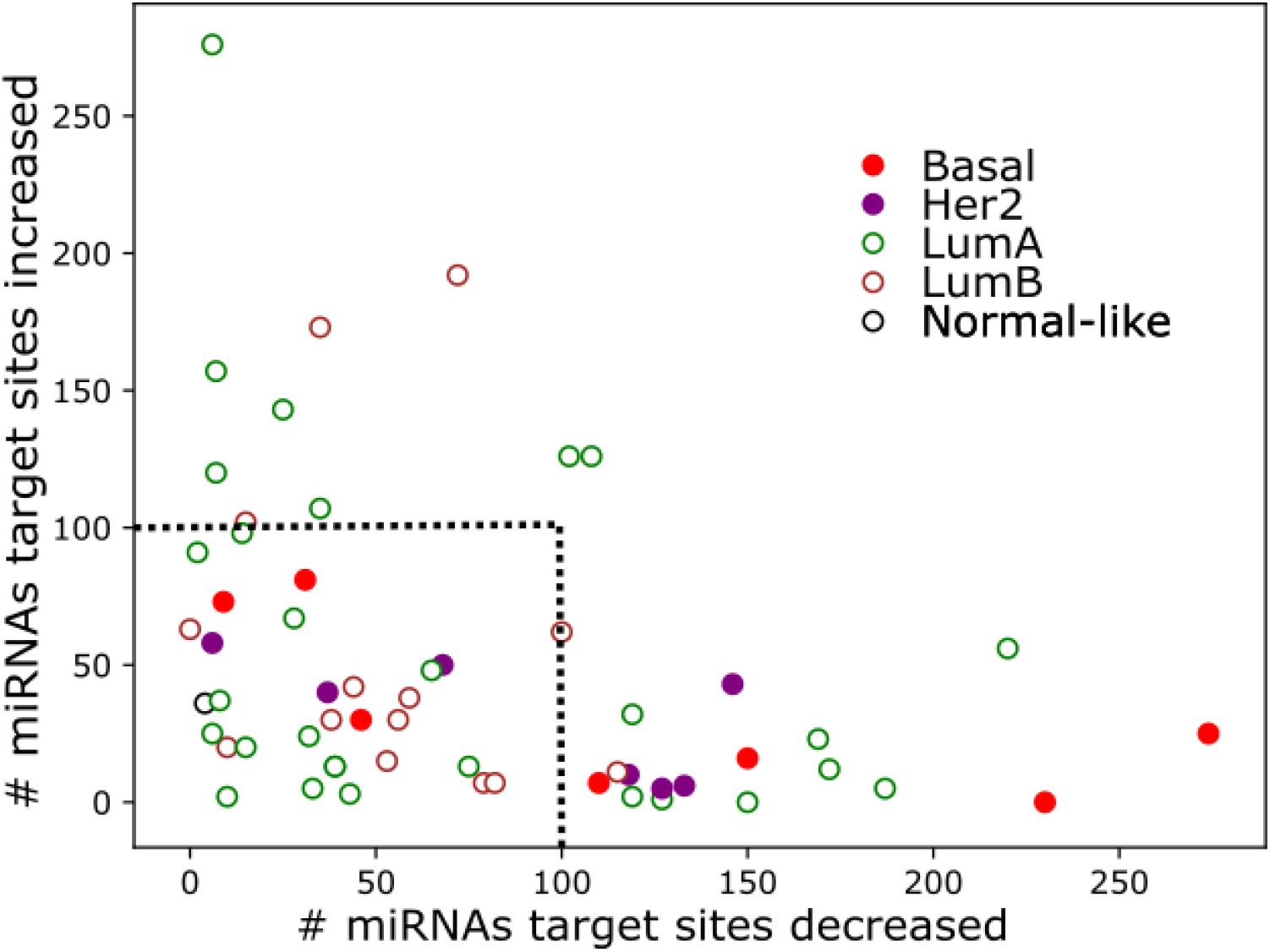
Number of miRNAs of which binding sites are increased (y-axis) or decreased (x-axis) for each tumor-normal sample colored by the breast tumor subtype. The black dotted rectangle represents not-high modifications.

**S. Figure 3.**
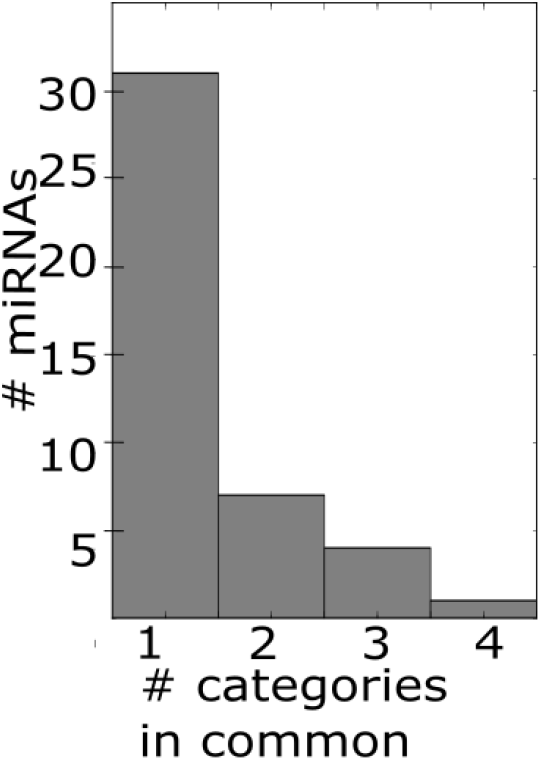
Number of miRNAs validated for breast tumor progression and treatment against how many validation category they are in.

**S. Figure 4.**
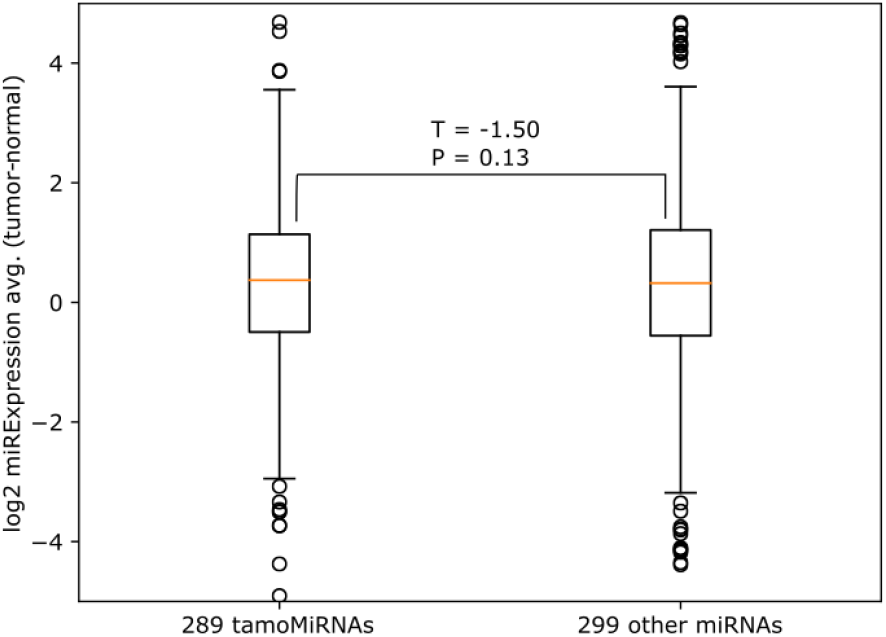
Expression difference (log 2, tumor vs. normal) of tamoMiRNAs and other miRNAs.

**S. Figure 5.**
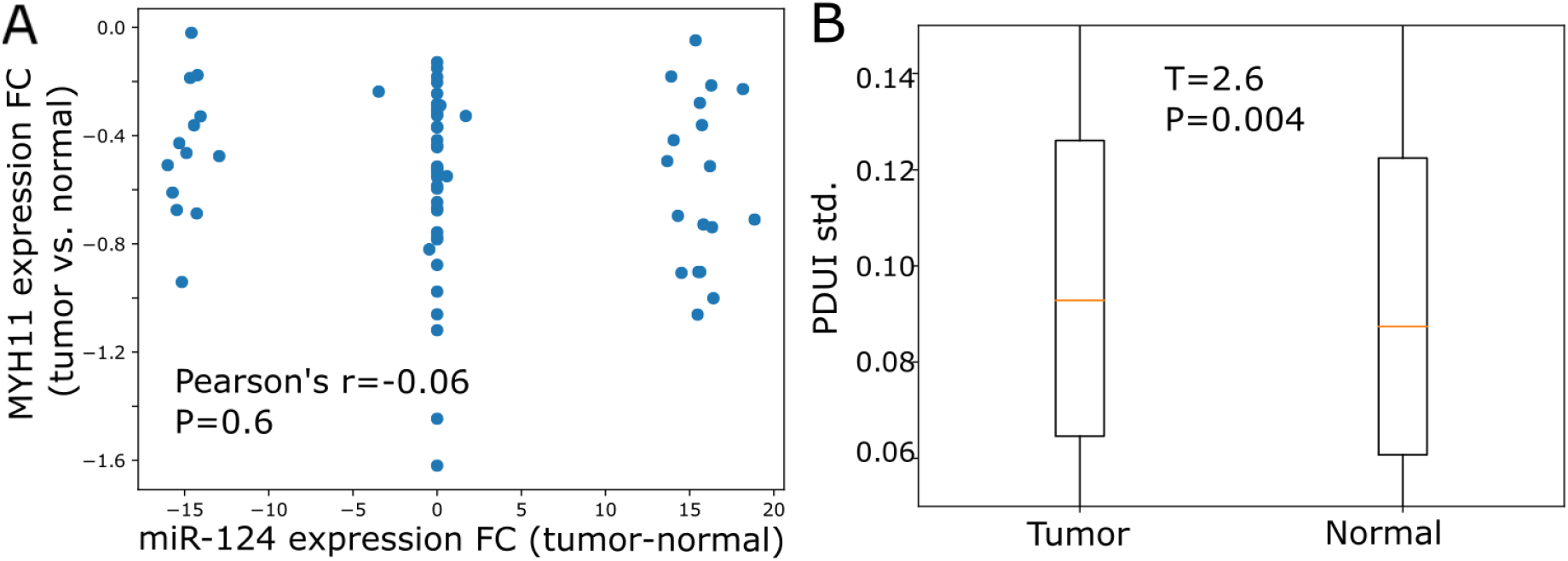
**A.** Scatterplot of MYH11 expression fold change values against expression fold change values of the mediating miRNA (miR-124/124ab/506). **B.** PDUI standard deviation o 2,862 genes with 3’-UTR usage status in tumor and normal whose PDUI values were available for all 70 tumor-normal pairs.

**Supplementary Table 1.** List of miRNAs with the number of patients in which APA modifies the target sites with the indication of whether the miRNA is found for each category of miRNAs with clinical potential.

**Supplementary Table 2.** List of miRNAs with the values for *MiRs_PDUI_n_*(*miR_j_*), *MiRs_PDUI_t_*(*miR_j_*), *MiRs_n_*(*miR_j_*), and *MiRs_t_*(*miR_j_*) followed by χ^2^ value and PRIMATA-APA call.

**Supplementary Table 3.** MiEAA analysis result for 99 tamoMiRs and 109 other miRNAs.

**Supplementary Table 4.** Ingenuity Pathway Analysis report on tamoMiRNA target genes.

## References

[1] M. Chen et al., “3 ‘ UTR lengthening as a novel mechanism in regulating cellular senescence,” pp. 285–294, 2018.

[2] G. P. Dimri et al., “A biomarker that identifies senescent human cells in culture and in aging skin in vivo.,” Proc. Natl. Acad. Sci. U. S. A., vol. 92, no. 20, pp. 9363–7, 1995.

[3] R. A. Busuttil, M. Rubio, M. E. T. Dollé, J. Campisi, and J. Vijg, “Oxygen accelerates the accumulation of mutations during the senescence and immortalization of murine cells in culture.,” Aging Cell, vol. 2, no. 6, pp. 287–294, 2003.

[4] C. López-Otín, M. A. Blasco, L. Partridge, M. Serrano, and G. Kroemer, “The hallmarks of aging,” Cell, vol. 153, no. 6, 2013.

[5] D. Muñoz-Espín and M. Serrano, “Cellular senescence: From physiology to pathology,” Nat. Rev. Mol. Cell Biol., vol. 15, no. 7, pp. 482–496, 2014.

[6] Z. Xia et al., “Dynamic Analyses of Alternative Polyadenylation from RNA-Seq Reveal Landscape of 3 ‘ UTR Usage Across 7 Tumor Types,” Nat. Commun., pp. 1–38, 2014.

[7] Y. Xiang et al., “Comprehensive Characterization of Alternative Polyadenylation in Human Cancer,” vol. 110, no. November 2017, pp. 1–11, 2018.

[8] H. J. Park et al., “3’ UTR shortening represses tumor-suppressor genes in trans by disrupting ceRNA crosstalk,” Nat. Genet., vol. 50, pp. 783–789, 2018.

[9] H. J. Park, S. Kim, and W. Li, “Model-based analysis of competing-endogenous pathways (MACPath) in human cancers,” PLoS Comput. Biol., vol. 22, no. 14, 2018.

[10] F. Zhenjiang, S. Kim, B. Diergaarde, and H. J. Park, “3’-UTR shortening disrupts ceRNA crosstalk of housekeeping genes resulting in subtype-specific breast cancer development,” bioRxiv, p. 601526, Jan. 2019.

[11] U. Ala et al., “Integrated transcriptional and competitive endogenous RNA networks are cross-regulated in permissive molecular environments.,” Proc. Natl. Acad. Sci. U. S. A., vol. 110, no. 18, pp. 7154–9, Apr. 2013.

[12] A. L. Sarver and S. Subramanian, “Competing endogenous RNA database.,” Bioinformation, vol. 8, no. 15, pp. 731–3, Jan. 2012.

[13] T. C. G. A. Network, “Comprehensive molecular portraits of human breast tumours.,” Nature, vol. 490, no. 7418, pp. 61–70, Oct. 2012.

[14] K. C. H. Ha, B. J. Blencowe, and Q. Morris, “QAPA: a new method for the systematic analysis of alternative polyadenylation from RNA-seq data,” pp. 1–18, 2018.

[15] C. Ye, Y. Long, G. Ji, Q. Q. Li, and X. Wu, “Genome analysis APAtrap: identification and quantification of alternative polyadenylation sites from RNA-seq data,” vol. 34, no. January, pp. 1841–1849, 2018.

[16] Z. Huang and E. C. Teeling, “ExUTR: A novel pipeline for large-scale prediction of 3’-UTR sequences from NGS data,” BMC Genomics, vol. 18, no. 1, pp. 1–11, 2017.

[17] W. Wang, Z. Wei, and H. Li, “A change-point model for identifying 3’UTR switching by next-generation RNA sequencing,” Bioinformatics, vol. 30, no. 15, pp. 2162–2170, 2014.

[18] L. Salmena, L. Poliseno, Y. Tay, L. Kats, and P. P. Pandolfi, “A ceRNA hypothesis: the Rosetta Stone of a hidden RNA language?,” Cell, vol. 146, no. 3, pp. 353–8, Aug. 2011.

[19] D. P. Bartel, “MicroRNAs: target recognition and regulatory functions.,” Cell, vol. 136, no. 2, pp. 215–33, Jan. 2009.

[20] T. Cancer Genome Atlas Network, “Comprehensive molecular portraits of human breast tumours,” 2012.

[21] X. Dai et al., “Breast cancer intrinsic subtype classification, clinical use and future trends,” Am J Cancer Res, vol. 5, no. 10, pp. 2929–2943, 2015.

[22] X. Dai, H. Cheng, Z. Bai, and J. Li, “Breast cancer cell line classification and Its relevance with breast tumor subtyping,” J. Cancer, vol. 8, no. 16, pp. 3131–3141, 2017.

[23] Y. Peng and C. M. Croce, “The role of MicroRNAs in human cancer,” Signal Transduct. Target. Ther., vol. 1, no. November 2015, p. 15004, 2016.

[24] S. Kurozumi, Y. Yamaguchi, M. Kurosumi, M. Ohira, and H. Matsumoto, “Recent trends in microRNA research into breast cancer with particular focus on the associations between microRNAs and intrinsic subtypes,” vol. 62, no. 1, pp. 15–24, 2016.

[25] A. Mehta and D. Baltimore, “MicroRNAs as regulatory elements in immune system logic,” Nat. Rev. Immunol., vol. 16, no. 5, pp. 279–294, 2016.

[26] E. van Schooneveld, H. Wildiers, I. Vergote, P. B. Vermeulen, L. Y. Dirix, and S. J. Van Laere, “Dysregulation of microRNAs in breast cancer and their potential role as prognostic and predictive biomarkers in patient management,” Breast Cancer Res., vol. 17, no. 1, p. 21, 2015.

[27] K. S. Pollard, M. J. Hubisz, K. R. Rosenbloom, and A. Siepel, “Detection of nonneutral substitution rates on mammalian phylogenies,” Genome Res., vol. 20, no. 1, pp. 110–121, 2010.

[28] A. Kozomara and S. Griffiths-Jones, “miRBase: annotating high confidence microRNAs using deep sequencing data.,” Nucleic Acids Res., vol. 42, no. Database issue, pp. D68–73, Jan. 2014.

[29] C. P. Masamha et al., “CFIm25 links alternative polyadenylation to glioblastoma tumour suppression.,” Nature, vol. 510, no. 7505, pp. 412–416, May 2014.

[30] C. Backes, Q. T. Khaleeq, E. Meese, and A. Keller, “MiEAA: MicroRNA enrichment analysis and annotation,” Nucleic Acids Res., vol. 44, no. W1, pp. W110–W116, 2016.

[31] C. H. Ooi et al., “A densely interconnected genome-wide network of micrornas and oncogenic pathways revealed using gene expression signatures,” PLoS Genet., vol. 7, no. 12, 2011.

[32] K. J. Mavrakis et al., “A cooperative microRNA-tumor suppressor gene network in acute T-cell lymphoblastic leukemia (T-ALL),” Nat. Genet., vol. 43, no. 7, pp. 673–678, 2011.

[33] Y. Hashimoto, Y. Akiyama, and Y. Yuasa, “Multiple-to-Multiple Relationships between MicroRNAs and Target Genes in Gastric Cancer,” PLoS One, vol. 8, no. 5, 2013.

[34] F. Sanchez-Vega et al., “Oncogenic Signaling Pathways in The Cancer Genome Atlas,” Cell, vol. 173, no. 2, pp. 321–337.e10, 2018.

[35] S. P. Nana-Sinkam and C. M. Croce, “MicroRNA regulation of tumorigenesis, cancer progression and interpatient heterogeneity: Towards clinical use,” Genome Biol., vol. 15, no. 9, pp. 1–9, 2014.

[36] N. Vickaryous et al., “Smooth-muscle myosin mutations in hereditary non-polyposis colorectal cancer syndrome,” Br. J. Cancer, vol. 99, no. 10, pp. 1726–1728, 2008.

[37] P. Liu et al., “Fusion Between Transcription Factor CBFß / PEBP2ß and a Myosin Heavy Chain in Acute Myeloid Leukemia Published by: American Association for the Advancement of Science Stable URL: https://www.jstor.org/stable/2881859 JSTOR is a not-for-profit service tha,” vol. 261, no. 5124, pp. 1041–1044, 2019.

[38] S. Huang, Z. G. Gulzar, K. Salari, J. Lapointe, J. D. Brooks, and J. R. Pollack, “Recurrent deletion of CHD1 in prostate cancer with relevance to cell invasiveness,” Oncogene, vol. 31, no. 37, pp. 4164–4170, 2012.

[39] L. Burkhardt et al., “CHD1 Is a 5q21 tumor suppressor required for ERG rearrangement in prostate cancer,” Cancer Res., vol. 73, no. 9, pp. 2795–2805, 2013.

[40] P. Alhopuro et al., “Unregulated smooth-muscle myosin in human intestinal neoplasia,” Proc. Natl. Acad. Sci., vol. 105, no. 14, pp. 5513–5518, 2008.

[41] I. Gupta et al., “Alternative polyadenylation diversifies post-transcriptional regulation by selective RNA–protein interactions,” Mol. Syst. Biol., vol. 10, no. 719, p. 176, 2014.

[42] W. Chen, Q. Jia, Y. Song, H. Fu, G. Wei, and T. Ni, “Alternative Polyadenylation: Methods, Findings, and Impacts,” Genomics. Proteomics Bioinformatics, vol. 15, no. 5, pp. 287–300, Oct. 2017.

[43] Li. Song et al., “miR-494 suppresses the progression of breast cancer in vitro by targeting CXCR4 through the Wnt/β-catenin signaling pathway,” Oncol. Rep., vol. 34, no. 1, pp. 525–531, 2015.

[44] F. C. Sigloch, U. C. Burk, M. L. Biniossek, T. Brabletz, and O. Schilling, “miR-200c dampens cancer cell migration via regulation of protein kinase A subunits,” Oncotarget, vol. 6, no. 27, 2015.

[45] N. Hu et al., “MiR-520b regulates migration of breast cancer cells by targeting hepatitis B X-interacting protein and interleukin-8,” J. Biol. Chem., vol. 286, no. 15, pp. 13714–13722, 2011.

[46] T. Shalaby, G. Fiaschetti, M. Baumgartner, and M. A. Grotzer, “Significance and therapeutic value of miRNAs in embryonal neural tumors,” Molecules, vol. 19, no. 5, pp. 5821–5862, 2014.

[47] R. Gambari, E. Brognara, D. A. Spandidos, and E. Fabbri, “Targeting oncomiRNAs and mimicking tumor suppressor miRNAs: Ew trends in the development of miRNA therapeutic strategies in oncology (Review),” Int. J. Oncol., vol. 49, no. 1, pp. 5–32, 2016.

[48] Y. Xiang et al., “Comprehensive Characterization of Alternative Polyadenylation in Human Cancer,” JNCI J. Natl. Cancer Inst., vol. 110, no. March, pp. 1–11, 2017.

[49] D. Endesfelder et al., “Intratumor Heterogeneity and Branched Evolution Revealed by Multiregion Sequencing,” N. Engl. J. Med., vol. 366, no. 10, 2012.

[50] P. L. Bedard, A. R. Hansen, M. J. Ratain, and L. L. Siu, “Tumour heterogeneity in the clinic,” Nature, vol. 501, pp. 355–364, 2013.

[51] M. Goldman et al., “The UCSC Cancer Genomics Browser: update 2013.,” Nucleic Acids Res., vol. 41, no. Database issue, pp. D949–54, Jan. 2013.

[52] B. P. Lewis, C. B. Burge, and D. P. Bartel, “Conserved seed pairing, often flanked by adenosines, indicates that thousands of human genes are microRNA targets.,” Cell, vol. 120, no. 1, pp. 15–20, Jan. 2005.

[53] G. L. Papadopoulos, M. Reczko, V. a Simossis, P. Sethupathy, and A. G. Hatzigeorgiou, “The database of experimentally supported targets: a functional update of TarBase.,” Nucleic Acids Res., vol. 37, no. Database issue, pp. D155–8, Jan. 2009.

[54] F. Xiao, Z. Zuo, G. Cai, S. Kang, X. Gao, and T. Li, “miRecords: An integrated resource for microRNA-target interactions,” Nucleic Acids Res., vol. 37, no. November 2008, pp. 105–110, 2009.

[55] S.-D. Hsu et al., “miRTarBase update 2014: an information resource for experimentally validated miRNA-target interactions.,” Nucleic Acids Res., vol. 42, no. Database issue, pp. D78–85, Jan. 2014.

[56] H. Dvinge et al., “The shaping and functional consequences of the microRNA landscape in breast cancer.,” Nature, vol. 497, no. 7449, pp. 378–82, May 2013.

[57] M. P. Hamilton et al., “Identification of a pan-cancer oncogenic microRNA superfamily anchored by a central core seed motif.,” Nat. Commun., vol. 4, p. 2730, Jan. 2013.

[58] M. G. G’Sell, S. Wager, A. Chouldechova, and R. Tibshirani, “Sequential Selection Procedures and False Discovery Rate Control,” arXiv Prepr., p. 27, 2013.

[59] M. Mancini and A. Toker, “NFAT proteins: Emerging roles in cancer progression,” Nat. Rev. Cancer, vol. 9, no. 11, pp. 810–820, 2009.

[60] G. C. Harburg and L. Hinck, “Navigating breast cancer: Axon guidance molecules as breast cancer tumor suppressors and oncogenes,” J. Mammary Gland Biol. Neoplasia, vol. 16, no. 3, pp. 257–270, 2011.

[61] C. Corzo et al., “The MYC oncogene in breast cancer progression: From benign epithelium to invasive carcinoma,” Cancer Genet. Cytogenet., vol. 165, no. 2, pp. 151–156, 2006.

[62] J. Cai, “Decreased Pigment Epithelium-Derived Factor Expression in Human Breast Cancer Progression,” Clin. Cancer Res., vol. 12, no. 11, pp. 3510–3517, 2006.

[63] M. L. Nierodzik and S. Karpatkin, “Thrombin induces tumor growth, metastasis, and angiogenesis: Evidence for a thrombin-regulated dormant tumor phenotype,” Cancer Cell, vol. 10, no. 5, pp. 355–362, 2006.

[64] K. K. Chan et al., “Erythropoietin drives breast cancer progression by activation of its receptor EPOR,” Oncotarget, vol. 8, no. 24, pp. 38251–38263, 2015.

[65] Z. Li et al., “miR-29a regulated ER-positive breast cancer cell growth and invasion and is involved in the insulin signaling pathway,” Oncotarget, vol. 8, no. 20, pp. 32566–32575, 2017.

[66] F. J. Sulzmaier, C. Jean, and D. D. Schlaepfer, “FAK in cancer: Mechanistic findings and clinical applications,” Nat. Rev. Cancer, vol. 14, no. 9, pp. 598–610, 2014.

[67] K. Banerjee and H. Resat, “Constitutive activation of STAT3 in breast cancer cells: A review,” Int. J. Cancer, vol. 138, no. 11, pp. 2570–2578, 2016.

